# Status epilepticus induces chronic silencing of burster and dominance of regular firing neurons during sharp wave-ripples in the mouse subiculum

**DOI:** 10.1101/2022.05.13.491697

**Authors:** Kristina Lippmann, Zin-Juan Klaft, Seda Salar, Jan-Oliver Hollnagel, Anna Maslarova

## Abstract

Sharp wave-ripples (SWRs) are hippocampal oscillations associated with memory consolidation. The subiculum, as the hippocampal output structure, ensures that hippocampal memory representations are transferred correctly to the consolidating neocortical regions. Because patients with temporal lobe epilepsy often develop memory deficits, we hypothesized that epileptic networks may disrupt subicular SWRs. We therefore investigated the impact of experimentally induced status epilepticus (SE) on subicular SWRs and contributing pyramidal neurons using electrophysiological recordings in mouse hippocampal slices. Subicular SWRs expressed hyperexcitable features post-SE, including increased ripple and unit activity. While regular firing neurons normally remain silent during SWRs, selective disinhibition recruited more regular firing neurons for action potential generation during SWRs post-SE. By contrast, burster neurons generated fewer action potential bursts during SWRs post-SE. Furthermore, altered timing of postsynaptic and action potentials suggested distorted neuronal recruitment during SWRs. Distorted subicular SWRs may therefore impair information processing and memory consolidation in epilepsy.

## Introduction

Sharp wave-ripples (SWRs) are transient high-frequency oscillations emerging during rest and slow-wave sleep in the hippocampus (Buzsáki, 2015; Shein-Idelson et al., 2016; Ylinen et al., 1995). They are believed to contribute to memory consolidation by providing a temporal framework in which participating neurons generate sequences of action potentials (APs) that represent a temporally compressed replay of firing series, previously generated during learning or exploratory behavior (Colgin, 2016; Diba and Buzsáki, 2007; Wilson and McNaughton, 1994). SWRs are thought to assist in transferring hippocampal representations to cortical structures for memory consolidation (Buzsáki, 2015; Chrobak and Buzsáki, 1996; Nitzan et al., 2020; Siapas and Wilson, 1998). The subiculum, the output structure of the hippocampus, is involved in spatial learning and memory (Barry et al., 2006; Burgess and O’Keefe, 1996; Norimoto et al., 2013; O’Keefe and Burgess, 1996), and was recently shown to be a generator for a subfraction of SWRs itself (Imbrosci et al., 2021; Norimoto et al., 2013). Intact subicular function is therefore essential to ensure that hippocampal representations are correctly generated, modified and transferred to the consolidating cortical areas.

This “exit gatekeeper” function is also relevant in temporal lobe epilepsy involving the hippocampal formation. Here, the subiculum is commonly engaged in seizure generation and propagation (Benini and Avoli, 2005; Lévesque and Avoli, 2020; Toyoda et al., 2015, 2013). Since temporal lobe epilepsy is often associated with cognitive comorbidities like dysfunctions in learning and memory as well as in spatial orientation (Bell, 2012; Cimadevilla et al., 2014; Grewe et al., 2014; Helmstaedter and Elger, 2009), it is significant whether subicular SWRs are altered as a consequence of epileptic networks.

In the hippocampus of epileptic patients and rodents, fast (or ‘pathological’) ripples with frequencies between 150-600 Hz (Alvarado-Rojas et al., 2015; Bragin et al., 1999) have been observed in parallel to physiological SWRs (with ~150-250 Hz ripples; Draguhn et al., 2000). The discrimination of both types of ripple-like oscillations in the human subiculum can be challenging due to their similar shape and frequency (Alvarado-Rojas et al., 2015). Nevertheless, fast ripples were shown to occur in the vicinity of the seizure focus and are thus considered a marker for ictogenesis (Bragin et al., 1999; Jefferys et al., 2012; Jiruska et al., 2017). It was likewise observed that SWRs and interictal events could be discriminated by their amplitude and duration (Alvarado-Rojas et al., 2015; Karlócai et al., 2014). Moreover, increasing network excitability can lead to replacement of SWRs by interictal events, indicating that the neuronal network generating SWRs can switch to interictal or ictal activity, when a certain level of excitability is exceeded (Karlócai et al., 2014; Liotta et al., 2011). Whereas it is generally accepted that pathologic ripples, interictal events and physiological SWRs are distinct events, less is known about whether SWRs are altered in the epileptic hippocampus. Studies in kainic acid (kainate)-treated rats showed preserved SWRs in hippocampal area CA1 and in the entorhinal cortex (EC) (Bragin et al., 1999) with local field potential (LFP) characteristics similar to those in naïve animals. Yet, it remains unclear whether epileptogenic network reorganization affects the processing of SWRs in the subiculum that might result in modified modulation of the cortical target areas.

In the subiculum, two distinct types of pyramidal neurons with distinct downstream projections, regular firing (RF) and burster neurons, have been described (Graves et al., 2012; Greene and Totterdell, 1997; Kim and Spruston, 2012). Subicular SWRs seem to involve, besides interneurons, predominantly burster neurons, which are activated location-dependently during exploratory states and fire bursts of APs during SWRs. AP bursts were shown to better encode spatial information compared to single spikes (Simonnet and Brecht, 2019) and burster neurons projecting to the EC and prefrontal cortex are crucial for propagating SWRs to extrahippocampal storage pathways (Böhm et al., 2018). Burster neurons can undergo modifications in epilepsy and gain additional excitability by means of mismatch in ionic currents and decrease in inhibition (Chen et al., 2011; Jensen and Yaari, 1997). By contrast, RF neurons often remain silent during physiological SWRs (Böhm et al., 2015; Maslarova et al., 2015). However, additional excitation may alter the behavior of RF neurons and let them participate more in the SWR generation (Eller et al., 2015). We therefore hypothesized that the cellular involvement of subicular pyramidal neurons in SWRs may be altered by epilepsy-induced hyperexcitability.

To address this hypothesis we studied spontaneously occurring SWRs in acute, combined hippocampal-entorhinal cortex slices of mice (Maier et al., 2003) in an epilepsy model. To this end, we induced status epilepticus (SE) *in vivo* by kainate injection in the dorsal hippocampus and recorded SWRs in slices from the ventral hippocampus, remote from the ictogenic focus (Bragin et al., 1999; Cavalheiro et al., 1982; Dugladze et al., 2007). We observed that spontaneous SWR generation was preserved in the ventral subiculum of mice that underwent SE. However, SWRs expressed hyperexcitable features, such as more ripples and units, i.e., neuronal spikes. In addition, the overall percentage of SWRs with long first-to-last ripple durations, which have been particularly linked to memory consolidation (Fernández-Ruiz et al., 2019), was reduced. Post-SE SWRs recruited less burster neurons and more RF neurons compared to SWRs in naïve animals, which was accompanied by an increase and loss of inhibitory inputs onto burster and RF neurons, respectively. This differential cellular contribution may represent a disruption of fine-tuning and information encoding during SWR-associated memory processing.

## Results

### Kainate injection into the dorsal hippocampal CA1 reliably induces status epilepticus and chronic seizures

To study the effect of epileptogenesis on SWRs in subiculum, we chose the well-established mouse model of kainate-induced SE (Dugladze et al., 2007). To monitor chronic seizure development, we performed LFP recordings from the right area CA1 (rHippo) and electrocorticographic (ECoG) recordings from the right frontal lobe (rFront) and the left parietal lobe (lPariet, Fig. S1A) for 6-9 weeks after intrahippocampal kainate injections. To verify that electrode implantation did not itself induce seizures, we performed baseline recordings in the time range of one week to two hours before kainate injection. None of the animals (n = 0 of 9) presented epileptiform activity in any of the recorded brain regions (Fig. S1). Two weeks after recovery from electrode implantation, SE was induced by kainate injection into right CA1 area of the dorsal hippocampus and recordings were performed immediately afterwards. SE was characterized by recurrent generalized electrographic seizures visible in all recording sites (Fig. S1C, middle panels) as well as tonic-clonic motor seizures, and was induced in all injected animals. Kainate-induced SE reliably leads to spontaneous chronic seizures and histo-morphological changes in the hippocampus resembling epileptogenesis in humans with temporal lobe epilepsy (Bouilleret et al., 1999; Dugladze et al., 2007). To verify epileptogenesis in animals, we performed sporadic recordings for 1-2 hours once every 1-3 weeks, up to 9 weeks following SE. Indeed, we detected seizures in 67% of injected mice (n = 6 of 9). In the remaining three animals, seizures could have been missed due to rare and short recordings post-SE. In most animals, chronic seizures were detected initially within the first three weeks post-SE. Seizures typically evolved in the hippocampal region and propagated in all cases to the ipsilateral frontal cortex. With one exception (n = 5 of 6 mice with detected seizures) seizures also propagated to the contralateral hemisphere, as detected by the lPariet ECoG electrode (Fig. S1 C and D) implying that most seizures generalized in several brain regions.

### Subicular sharp waves are altered in post-SE animals

We next aimed to compare subicular SWRs between control and post-SE animals in acute horizontal slices from ventral hippocampi of both cohorts. First, we performed simultaneous LFP recordings from the hippocampal areas CA3, CA1 and the subiculum in order to confirm the origin of SWRs (Fig. 1A). As previously described (Maier et al., 2003; Maslarova et al., 2015; Wu et al., 2006), SWRs propagated from CA3 to CA1 and subsequently to the subiculum in naïve mice (Fig. 1B, black traces). The propagation was preserved in animals that underwent SE (Fig. 1B, blue traces). For further quantification, we recorded a large number of subicular SWRs (control: 15463 events from 15 slices of 10 animals, post-SE: 19094 events from 24 slices of 7 animals) and conducted specific analyses of the different components of a SWR, namely the sharp wave (SPW), i.e., the slow wave component of the SWR, the ripples and units, i.e., single neuron firing. For analyzing SPWs, the data was filtered with an 80 Hz low-pass filter (Fig. 1C). SPWs occurred at a median rate of 106 (interquartile range (IQR): 80) per minute in control, whereas in post-SE animals, the SPW rate was significantly lower with 69 (89) SPWs per minute (Kolmogorov-Smirnov test (KS) p-value (p) = 0.0045, Fig. 1D). SPW duration was similar between groups (control: 25.6 (14.7) ms vs. post-SE (30.8 (23.5) ms, Fig. 1E), however, in post-SE animals SPW amplitudes were larger (control: 0.095 (0.119) mV vs. post-SE: 0.100 (0.059) mV, KS p = 0.003, Fig. 1F). In short, the propagation of SWRs from CA3 to the subiculum was conserved, while the subicular SPW rate was reduced and amplitudes were increased after SE.

**Figure 1.**
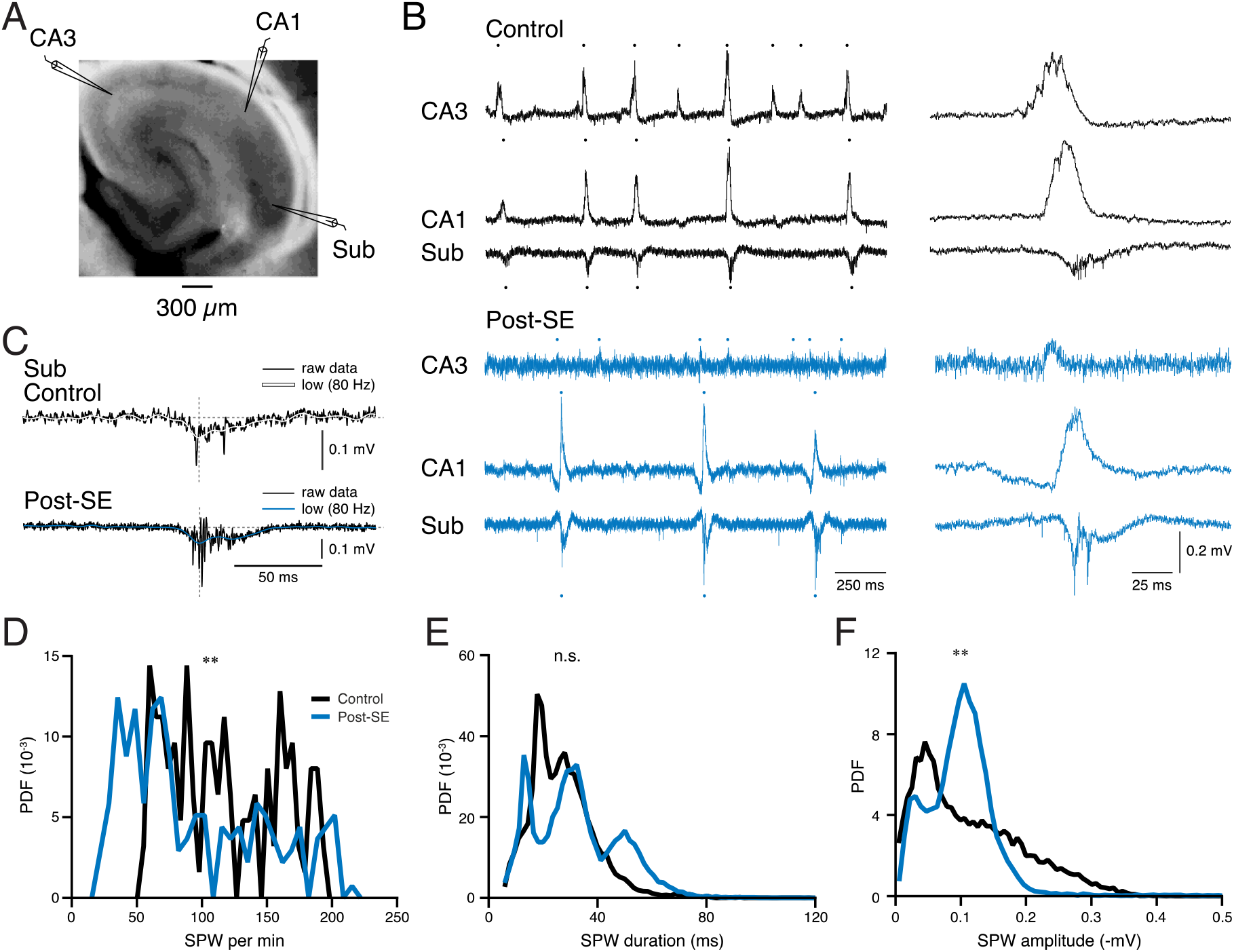
Altered subicular sharp waves (SPWs) post status epilepticus (post-SE). **(A)** Photograph of a horizontal ventral hippocampal slice. Local field potential (LFP) recording sites of CA3, CA1 and subiculum (Sub) with schematic electrodes are depicted. **(B)** Left: Spontaneously occurring SWRs in acute slices from a control (black traces) and a post-SE mouse (blue traces). Dots above (for CA3 and CA1) or below (for subiculum) traces depict identified SWRs. Right: Zoom-ins on the fifth (control) and second (post-SE) event. **(C)** Example SWRs from control (top) and post-SE (bottom) with the lowpass (‘low’) filtered trace overlaid (smooth white and blue line, respectively). Lowpass data were used for analysis of SPW frequencies, amplitudes and durations. The dotted horizontal and vertical lines depict the baseline and the trough of the SPW, respectively. **(D-F)** Comparison of SPW events between control (total events n = 15463 from 15 slices and 10 animals) and post-SE animals (n = 19094 from 24 slices and 7 animals) regarding the probability density function of the SPW rate (D), duration (E), and amplitude (F). ** p < 0.01.

### Subicular SWRs from post-SE mice contain more and larger ripples and units

Thereafter, we investigated the fast components of SWR oscillations – namely the ripples, which are believed to represent interneuron-driven synchronizations of pyramidal neurons during SWRs (Buzsáki, 2015; Csicsvari et al., 1999; Ylinen et al., 1995) and are essential for carrying information content (Fernández-Ruiz et al., 2019). Besides, we assessed potential alterations of units as correlates of contributing neuronal spikes. In both animal groups, bandpass filtering of the raw data (120-300 Hz and 600-4000 Hz, respectively) revealed oscillations in the ripple frequency range (Draguhn et al., 2000; Eller et al., 2015; Imbrosci et al., 2021; Wu et al., 2006) and single-unit activity superimposed on the slow negative field potential of subicular SPWs (Fig. 2A and B). The number of ripples per SWR was significantly higher in post-SE mice (KS p = 2.4e^-5^, Fig. 2C), with yet similar ripple frequencies in both animal groups (control: 200 (77) Hz vs. post-SE: 208 (74) Hz, Fig. 2E), but larger ripple amplitudes in post-SE animals (control: 42 (30) μV vs. post-SE: 53 (40) μV, KS p = 2.4e^-5^, Fig. 2F). Having observed a difference in the ripple count per SWR, it was surprising that the overall first-to-last ripple duration (‘ripple duration’) did not differ between the two groups (control: 12.4 (23) ms vs. post-SE: 13.0 (17) ms, KS p = 0.26, Fig. 2G). Because long-duration ripples have been linked to better memory performance (Fernandez Ruiz 2019), we assessed the proportion of SWRs with a first-to-last ripple duration exceeding 100 ms. Indeed, in post-SE animals we found a significantly lower proportion of such, long-ripple-duration SWRs’ (control: 4.1% (249/6026 SWRs) vs. post-SE: 2.7% (296/11061 SWRs), Chi-square p < 0.001, Fig. 2G inset, H).

**Figure 2.**
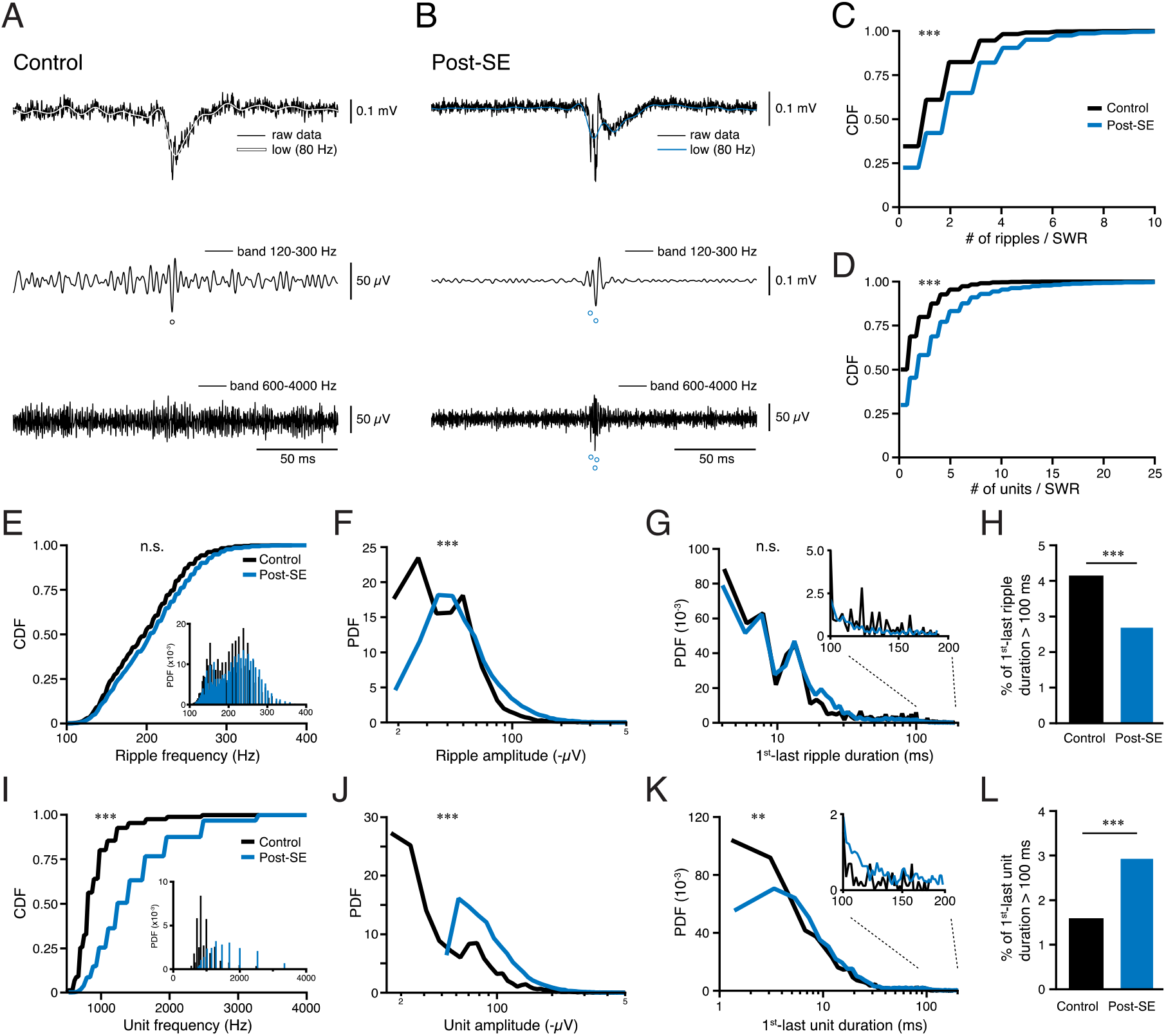
Subicular SWRs contain more and larger ripples and units post-SE. **(A-B)** Representative traces of a control (A) and post-SE (B) SWR. Raw data (top black traces) were lowpass filtered (80 Hz, top white and blue traces) for analysis of SPW parameters (cf. Fig. 1) and bandpass filtered for quantifying properties of ripples (120 - 300 Hz, middle) and units (600 - 4000 Hz, bottom traces). Circles denote ripples or units detected in the respective bandpass filtered traces. **(C)** Normalized cumulative density function (CDF) of the number of ripples per SWRs compared between control (n = 20171) and post-SE (n = 39808). **: p < 0.01, ***: p < 0.001. **(D)** As in C but for the number of units per SWRs (control n = 20338, post-SE n = 56738). **(E)** CDF of the ripple frequencies with the inset showing the PDF of the data. (**F**) PDF of the ripple amplitudes. **(G)** PDF of the duration between the first and the last ripple (‘ripple duration’) for SWRs containing at least two ripples (n = 6026 and 11061 in control and post-SE, respectively). Zoom-in displays the data range for ripple durations > 100 ms. **(H)** Percentage of 1^st^-last ripple durations > 100 ms in control and post-SE SWRs. **(I-L)** Same analysis for units as described above for ripples (E-H; n = 4810 and 10441 in control and post-SE, respectively).

To evaluate neuronal firing during SWRs, we analyzed unit activity. In post-SE animals, we observed a significantly higher number of units per SWR (KS p = 9.6e^-18^, Fig. 2D). Additionally, the unit frequency (control: 833 (286) Hz vs. post-SE: 1250 (667) Hz, KS p = 3.2e^-4^, Fig. 2I) and amplitude (control: 46 (47) μV vs. post-SE: 88 (49) μV, KS p = 2.4e^-5^, Fig. 2J) were significantly increased in post-SE mice. In accordance with the higher number of units per SWR, the first-to-last unit duration was significantly prolonged in post-SE mice (control: 6.4 (13) ms vs. post-SE: 9.0 (16), KS p = 0.003, Fig. 2K). This was also true for the proportion of SWRs with a first-to-last unit duration exceeding 100 ms and stood in contrast to the ripple durations (“long-unit-duration SWRs”, control: 1.6% (75/4810 SWRs) vs. post-SE: 2.9% (304/10441 SWRs), Chi-square p < 0.001, Fig. 2K inset, 2L), implying that some of the post-SE units emerged independently of the ripple activity. Increased numbers and amplitudes of ripples and units reveal an augmented cellular contribution during SWRs. However, the diminished first-to-last ripple durations but increased unit durations and frequencies indicate a distorted timing and, thus, information encoding of neuronal activity during subicular SWRs post-SE.

### Altered temporal dynamics in the sharp wave-ripple complex

To detect possible changes in the ripple/unit regulation, we investigated whether the time relations between ripples and units relative to the SPW trough were altered. It has previously been shown that epileptic ripples preceded SPWs in CA1 (Bragin et al., 1999). We found that the time relation of ripple trough to SPW trough was significantly different in control and post-SE mice, revealing a broader distribution of ripple timing and a later peak in post-SE mice (control: −3.4 (12) ms vs. post-SE: −2.6 (14), KS p = 9.9e^-15^, Fig. 3A). Moreover, the ripple buildup started earlier (−26 ms) in post-SE compared to control mice (−20 ms). In accordance with earlier evidence that interneuron-driven ripples phase-lock pyramidal cell (PC) firing (Chrobak and Buzsáki, 1996; Schlingloff et al., 2014), we observed, similar to ripples, a delayed peak of unit firing relative to the SPW trough (control: −0.6 (7) ms vs. post-SE: 0 (9), KS p = 0.001, Fig. 3B) and an earlier buildup of unit activity in post-SE animals. Moreover, unit activity revealed a second peak at −11 ms relative to the SPW trough in post-SE animals before transitioning back into the gaussian-like histogram at approximately −7 ms (Fig. 3B). We therefore analyzed events preceding the SWR trough by more than 7 ms, i.e., “early” events separately (Fig. 3, dashed boxes) and found that the proportion of “early” units had almost doubled in post-SE relative to control animals (control: 6.4% (1293/20338 units) vs. post-SE: 12.5% (7066/56738 units), Chi-square p < 0.001, Fig. 3B). The proportion of “early” ripples was likewise larger in post-SE mice (control: 15.6% (3151/20171 ripples) vs. post-SE: 16.5% (6575/39808 ripples), Chi-square p = 0.005, Fig. 3A).

**Figure 3.**
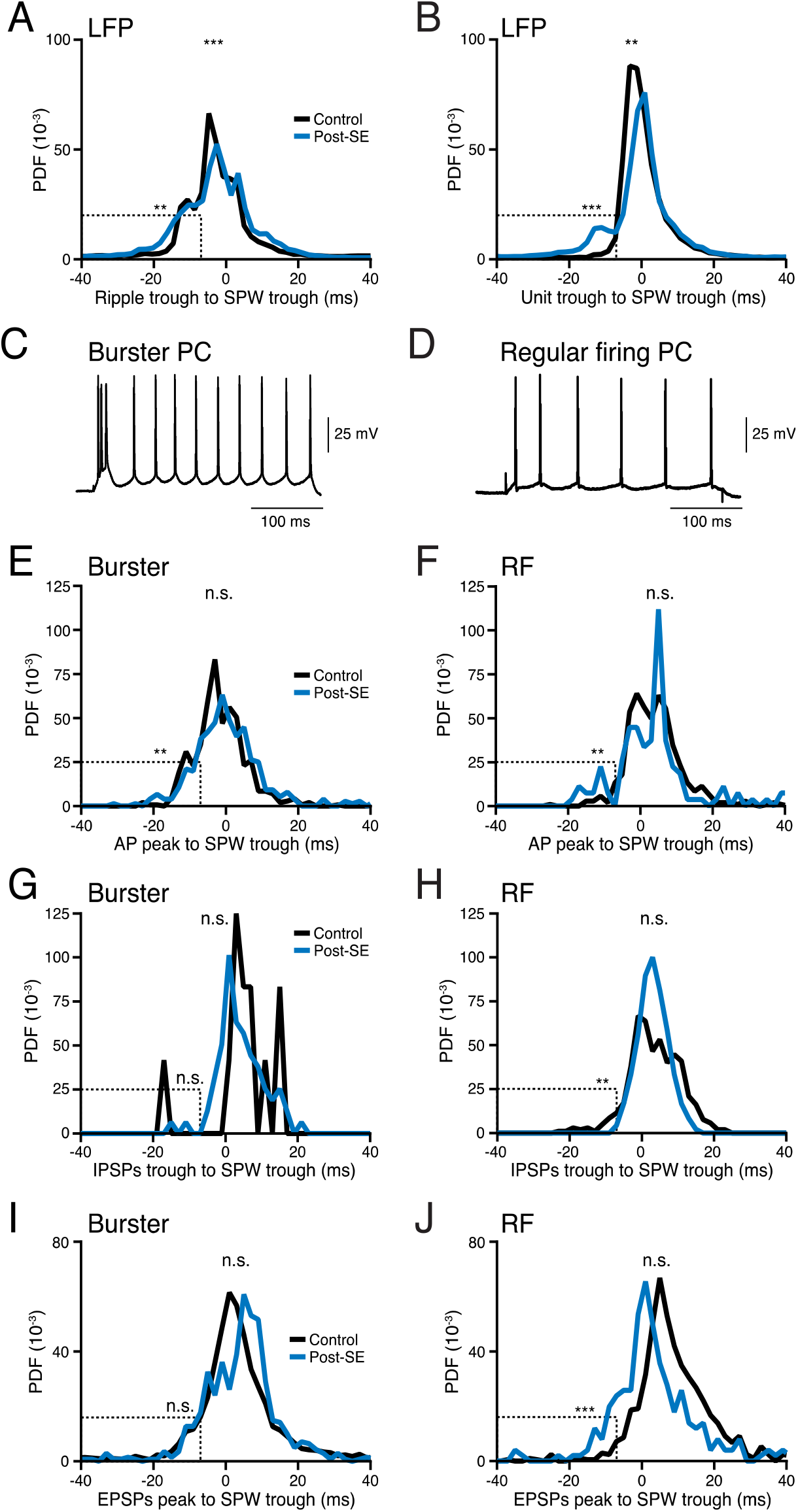
Altered temporal dynamics in the SWR-complex in post-SE mice. **(A)** PDF of the temporal relation of ripple troughs relative to the SWR trough (aligned to 0 ms) obtained using LFP recordings (same data sets as in Fig. 1–2). Dashed box indicates the proportion of ripples occurring at least 7 ms before the SWR trough. **(B)** Same analysis as in (A) for the temporal relation of unit troughs to the SWR trough. **(C-D)** Representative example traces of a burster and regular firing (RF) pyramidal cell (PC). Note that the burster PC generated an initial burst of APs with subsequent regular firing. **(E-F)** PDFs of the time of AP peak relative to the SWR trough of burster and RF neurons (E and F, respectively). Dashed boxes indicate the proportion of APs occurring at least 7 ms before the SWR trough. **(G-J)** PDFs of IPSP troughs (G, H) and EPSP peaks (I, J) relative to the corresponding SWR trough for burster (G and I) and RF neurons (H and J).

SWRs in the subiculum are generated by an interplay between interneurons and burster PCs, whereas the other subtype of principal neurons - regular firing (RF) PCs - are thought to remain silent (Böhm et al., 2015; Eller et al., 2015). To understand the cellular contributions to ripples and units and whether they change after SE, we performed sharp-electrode intracellular recordings from subicular neurons in parallel to LFP SWR recordings. Our recordings confirmed those two distinct types of subicular pyramidal neurons: burster and RF PCs (Fig. 3 C-D, for electrophysiological characterization see Fig. 4; Greene and Totterdell, 1997; Stewart, 1997; Taube, 1993). When comparing the temporal relation of their APs relative to the SPW trough, the AP histograms of burster neurons (Fig. 3E) correlated well with the ripple histograms (Fig. 3A) and the ones of RF neurons (Fig. 3F) with the unit histograms (Fig. 3B). As with the ripples, the histogram median of burster APs preceded the SPW trough by −2.3 (8.1) ms in control mice and was shifted to a slightly but insignificantly later time point in post-SE mice (−0.6 (10.2) ms, Fig. 3E). Interestingly, as in ripples, the proportion of “early” APs before-7 ms to the SPW trough was increased in burster neurons (control: 7.4% (53/714 APs) vs. post-SE: 12.4% (47/380 APs), Chi-square p = 0.007, dashed box in Fig. 3E). However, neither inhibitory nor excitatory postsynaptic potentials (IPSPs and EPSPs, respectively, Fig. 3G, 3I) revealed significant differences in their timing relative to the SPW trough. In RF neurons, APs, IPSPs and EPSPs from post-SE mice did not show a significant shift in the overall distributions, however they did show a significantly altered occurrences of “early events” with APs displaying an extra peak at ~11 ms preceding the SPW trough (proportion of “early” APs in control: 3.2% (17/532 APs) vs. post-SE: 9.0% (12/134 APs), Chi-square p = 0.0035, Fig. 3F). These data correlated well with the early unit peak in LFP recordings (Fig. 3B) and may also explain the higher proportion of long-unit-duration SWRs (Fig. 2K-L). Even though the median timing of the inhibitory inputs remained stable in both groups, there was a reduced proportion of early IPSPs in RFs (control: 4.0% (21/526 IPSPs) vs. post-SE: 0.4% (1/239 IPSPs), Chi-square p = 0.006, dashed box in Fig. 3H). By contrast, excitatory inputs occurred significantly earlier (proportion of EPSPs before −7 ms, control: 3.2% (34/1048 EPSPs) vs. post-SE: 11.8% (48/406 EPSPs), Chi-square p < 0.001, Fig. 3J). The reduced early inhibition and increased excitation may ultimately contribute to an increased AP firing up to −7 ms before the SPW trough, increasing temporal jitter of cellular contributions to SWRs and suggesting less precise memory consolidation.

**Figure 4.**
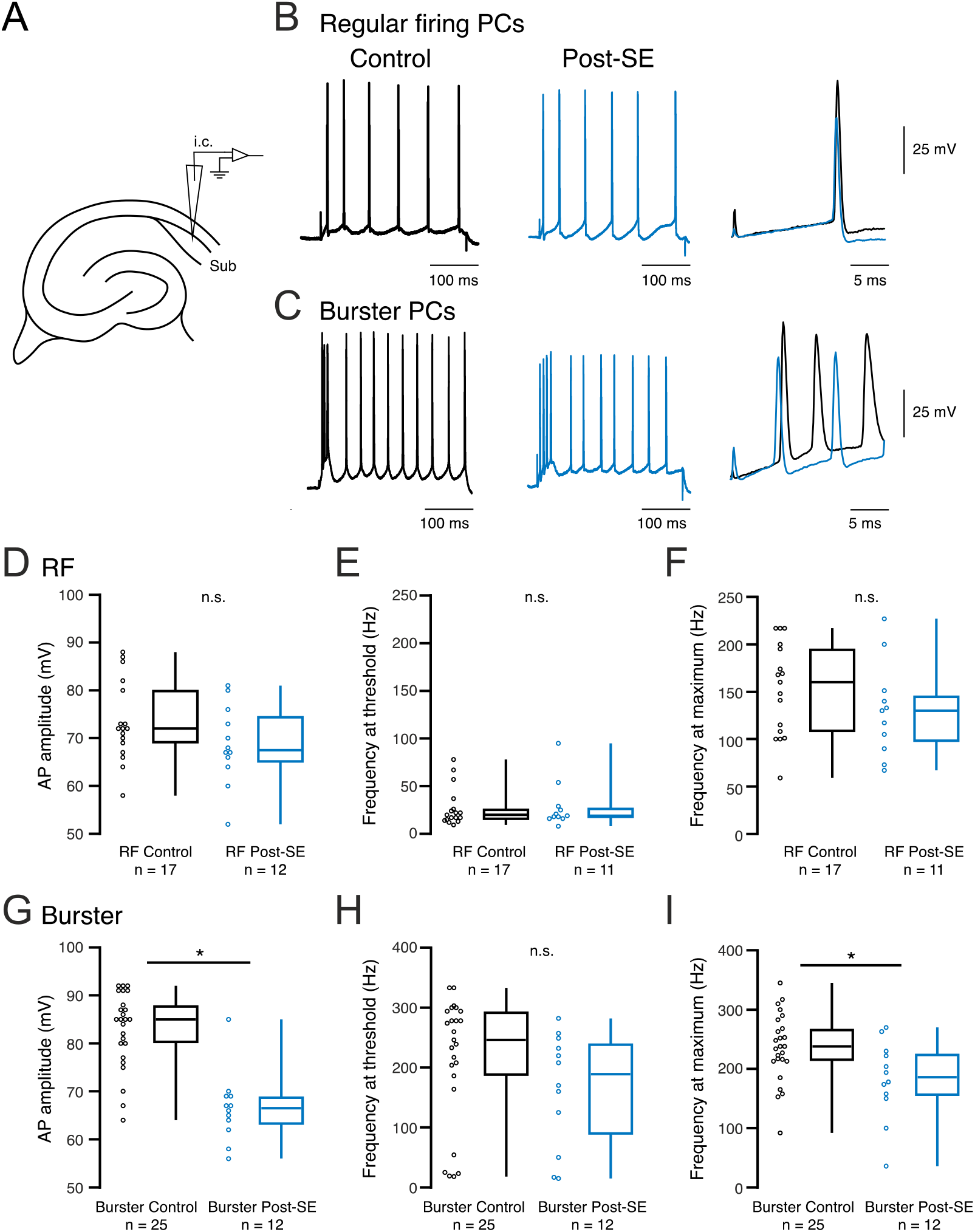
Subicular burster neurons undergo a decrease in intrinsic excitability upon status epilepticus. **(A)** Scheme of the sharp electrode recording site in the subiculum from horizontal ventral hippocampal slices. **(B-C)** Representative intracellular recordings of subicular neurons identified as regular firing (RF, B) and burster (C) pyramidal cells (PCs) from control and post-SE mice (black and blue traces, respectively). Responses to a depolarizing current step (amplitude set to twice the threshold amplitude) are displayed. Right panel: Zoom-in and overlay of the first action potentials (APs). **(D)** Amplitude of the first AP induced by a depolarizing step at the double threshold intensity of RF neurons. Dots represent single data points. Box plots show the median, Q25 and Q75. **(E)** Firing frequency of RF neurons calculated for the first two APs evoked by a depolarizing step at threshold current intensity. **(F)** Firing frequency at the chosen maximum current intensity (0.55 nA). **(G-I)** Same analysis for burster neurons. * p < 0.05; n denotes the number of cells per group.

### Only burster neurons undergo excitability changes under post-SE

Having identified higher numbers of ripples and units per SPW as well as altered AP and synaptic input timing in subicular SWRs in post-SE animals, we proceeded to investigate whether altered intrinsic properties of individual subicular RF and burster neurons (Stewart, 1997; Taube, 1993) underly these changes. In our intracellular recordings using sharp microelectrodes (Fig. 4A), both types of neurons, could be identified (see methods section *‘Electrophysiological characterization’*) in the control as well as in the post-SE group. While RF neurons fired at a fairly constant rate during depolarizing current steps (Fig. 4B), burster neurons generated a burst of APs at the beginning of a depolarizing step with subsequently more regular AP firing (Fig. 4C). Membrane properties and AP characteristics of RF neurons were similar in both groups (Table 1, Fig. 4D). Likewise, the firing rate was similarly low at threshold intensity (Fig. 4E) and increased with rising current intensity to an even extent to 160 (87) Hz in control animals and 130 (48) Hz in post-SE animals at maximum (max.) stimulation intensity (Fig. 4F). By contrast, in post-SE burster neurons the resting membrane potential (RMP) was slightly more positive (control neurons: −64.5 (4.3) mV, post SE neurons: −63.0 (5.5) mV, unpaired two-tailed student’s t-test (t-test) p = 0.031) and the AP amplitude (67 (6) mV) was significantly smaller than in control neurons (85 (8) mV, t-test p < 0.001, Fig. 4G). The maximum number of APs per burst was 4 (1) in control (n = 24) and 3.5 (3) in post-SE animals (n = 12). Although the firing frequency at threshold intensity did not differ significantly (Fig. 4H), the firing frequency at maximum stimulation intensity was reduced in post-SE animals (control: 238 (55) Hz, post-SE 186 (68) Hz, t-test p = 0.015, Fig. 4I). All in all, depolarizing step experiments revealed reduced amplitudes and firing frequencies in burster neurons.

**Table 1.**
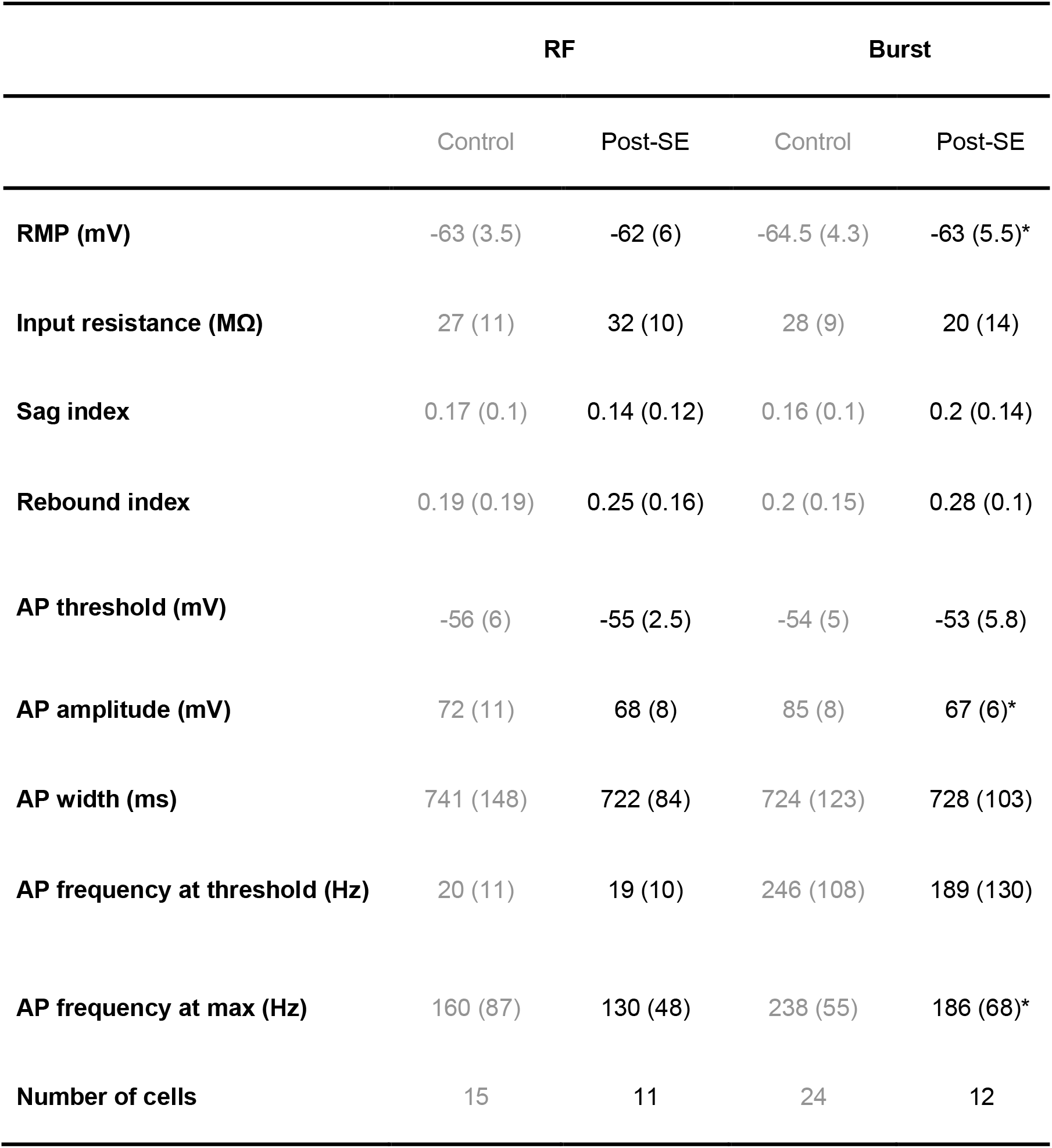
Intrinsic properties of stimulated subicular neurons. Intrinsic properties were evaluated during hyperpolarizing and depolarizing current steps of increasing intensity (70 pA step interval, 300 ms). All data are presented as median and, in brackets, IQR. RMP corresponds to the resting membrane potential. Sag and rebound indices were calculated from a 70 pA negative current step. AP amplitude and width (width at half-maximal amplitude) were measured at the double threshold current intensity. T-test: *p = 0.031, ** p < 0.001, *** p = 0.015.

|  | RF |  | Burst |  |
| --- | --- | --- | --- | --- |
|  | Control | Post-SE | Control | Post-SE |
| **RMP (mV)** | -63 (3.5) | -62 (6) | -64.5 (4.3) | -63 (5.5)* |
| **Input resistance (MΩ)** | 27 (11) | 32 (10) | 28 (9) | 20 (14) |
| **Sag index** | 0.17 (0.1) | 0.14 (0.12) | 0.16 (0.1) | 0.2 (0.14) |
| **Rebound index** | 0.19 (0.19) | 0.25 (0.16) | 0.2 (0.15) | 0.28 (0.1) |
| **AP threshold (mV)** | -56 (6) | -55 (2.5) | -54 (5) | -53 (5.8) |
| **AP amplitude (mV)** | 72 (11) | 68 (8) | 85 (8) | 67 (6)* |
| **AP width (ms)** | 741 (148) | 722 (84) | 724 (123) | 728 (103) |
| **AP frequency at threshold (Hz)** | 20 (11) | 19 (10) | 246 (108) | 189 (130) |
| **AP frequency at max (Hz)** | 160 (87) | 130 (48) | 238 (55) | 186 (68)* |
| **Number of cells** | 15 | 11 | 24 | 12 |

### Regular firing neurons receive less inhibitory drive and fire more APs during SWRs in post-SE mice

The reduced excitability in burster neurons did not provide an explanation for an increased number of ripples and units as well as their early occurrence in post-SE SWRs. However, this early occurrence was reminiscent of the earlier excitation and delayed inhibition in RF neurons. We therefore characterized the distribution of intracellular events during SWRs in subicular PCs in control and post-SE mice.

In a subset of SWR LFP recordings, we analyzed at least two minutes of intracellular data recorded in parallel to analyze cellular in- and outputs of the previously characterized RF and burster neurons during SWRs (Fig. 5A-B). In the majority of recorded neurons, each SWR was associated with at least one of three types of intracellular responses: APs, subthreshold EPSPs and IPSPs (see examples in Fig. 5B). While some neurons showed three types of activity (IPSPs, EPSPs and APs), others only showed two or one of them. Basic parameters of the intracellular events, such as amplitudes of all three types as well as AP width did not differ between control and post-SE animals (Table 2, Fig. 5C). However, we observed differences in the extent of inhibitory and excitatory drives onto RF neurons (Fig. 5D, 7). While in RFs from control mice IPSPs comprised 48% of all intracellular SWR-associated events, the proportion decreased to 18% in RFs from post-SE mice (Chi-square p < 0.001). Accordingly, post-SE RFs contained an increased proportion of EPSPs (control: 41% vs. post-SE: 63%, Chi-square p = 0.003). The total percentage of APs did not change significantly (control: 11% vs. post-SE: 19%, Chi-square p = 0.16). However, it seemed that the number of neurons receiving inhibitory drive was fundamentally reduced and the amount of neurons generating APs increased. To quantify these changes, we analyzed the proportion of neurons that received an excitatory or inhibitory input, or generated an AP at least once during SWRs (Fig. 5E). If more than one event type was detected in the same neuron, this neuron was counted for all occurring event groups. In accordance with previous reports from the subiculum (Böhm et al., 2015; Eller et al., 2015) reporting that RFs are usually silent during SWRs, 67% of the recorded RF neurons received IPSPs during SWRs. Strikingly, in post-SE mice, this was the case in only 18% of neurons (Chi-square p = 0.014). Even though the subthreshold excitatory input remained largely unchanged (EPSPs during SWRs in control: 67% vs. post-SE: 82%), the loss of inhibitory drive resulted in a significant increase of RF cells actively participating in SWRs. In fact, the proportion of RF cells firing APs during SWRs had more than doubled and comprised 73% (Chi-square p = 0.047) in post-SE mice, as opposed to merely 33% in control mice (Fig. 7, filling level of RF triangles). Altogether, in post-SE mice, more RF neurons were recruited in active firing during SWRs, since they had partially lost their inhibitory drive.

**Figure 5.**
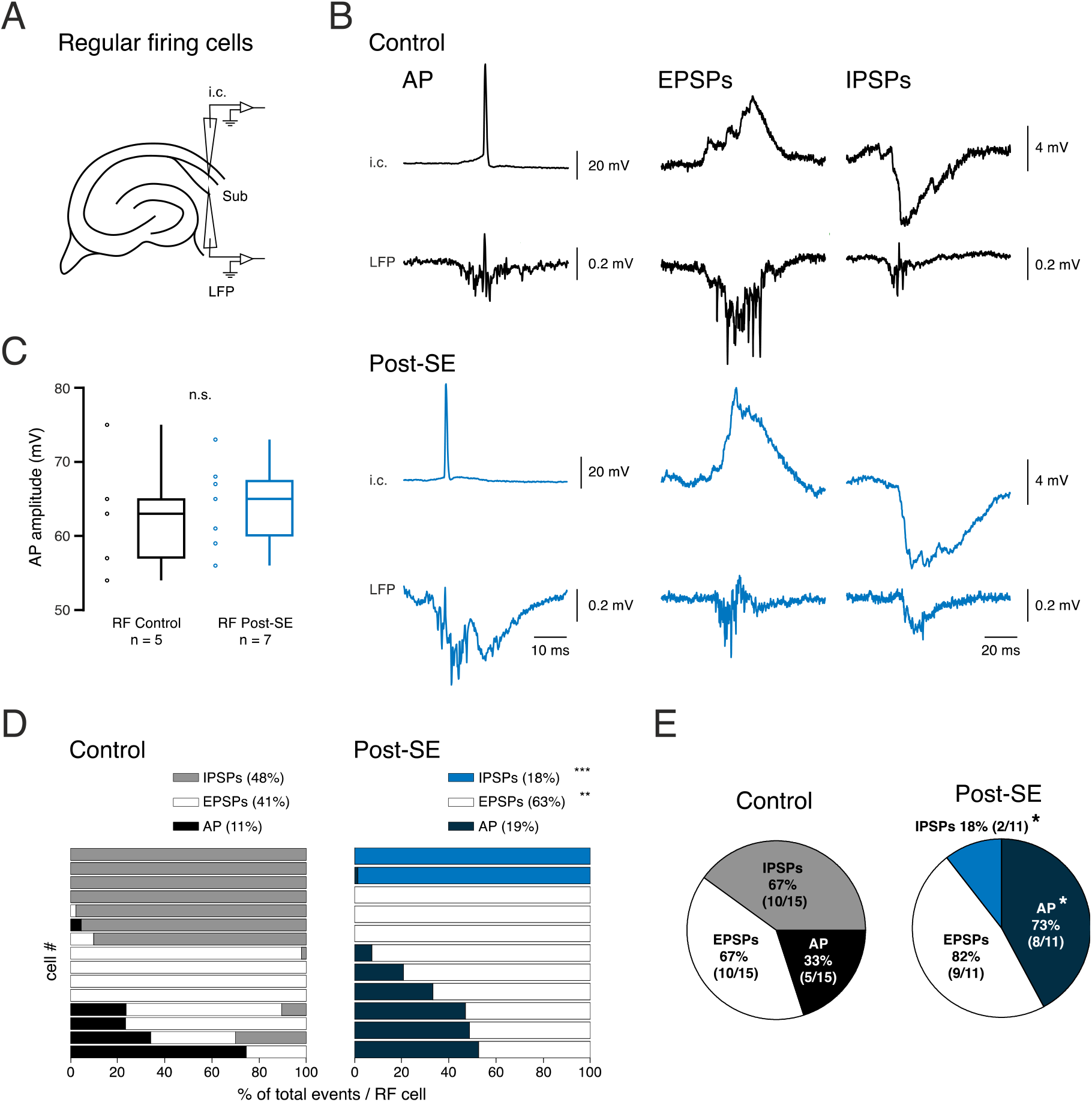
Increased activity of regular firing neurons in post-SE SWRs. **(A)** Scheme of the sharp electrode and LFP recording sides in the subiculum. **(B)** Representative traces of intracellular recordings (i.c., top traces) from a control (black traces) and post-SE (blue traces) regular firing subicular neuron illustrating the three types of intracellular activity observed during spontaneous SWRs (from left to right): action potentials (APs), excitatory and inhibitory postsynaptic potentials (EPSPs and IPSPs, respectively). Bottom traces depict simultaneous local field potential (LFP) recordings of subicular SWRs. **(C)** Amplitude of APs occurring in control and post-SE RF neurons during SWRs. **(D)** Distribution of the different types of SWR-coupled intracellular events for each recorded pyramidal cell (control n = 15 cells, post-SE n = 11 cells, same data set as in Fig. 3). Each horizontal bar represents one recorded neuron and 100% on the x axis refers to the total number of events for each recorded RF cell. Note that in some neurons SWRs are always associated with the same type of events, whereas in others a combination of events occurred. **(E)** Distribution of the % of neurons that can generate APs, IPSPs or EPSPs during SWRs. Note that neurons generating more than one type of event were added to all relevant event groups.

**Figure 6.**
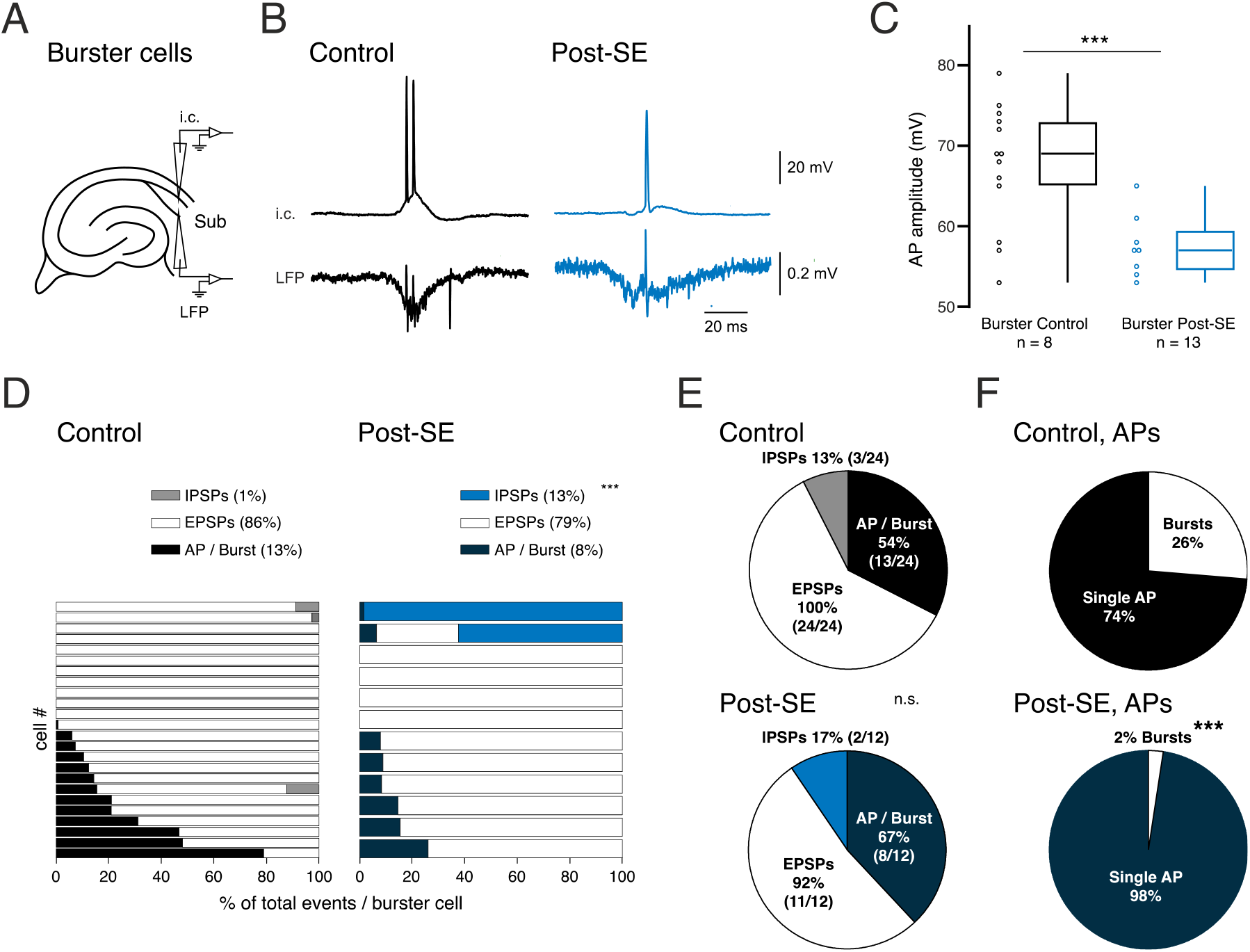
Reduced activity of burster neurons in post-SE SWRs. **(A)** Scheme of the sharp electrode and LFP recording sides in the subiculum. **(B)** Representative intracellular recordings from subicular burster neurons during spontaneous SWRs from control (black) and post-SE mice (blue trace), in which the cells fired a burst (upper traces, i.c.) during the SWR event (lower traces, LFP). **(C)** Amplitudes of APs occurring during SWRs. In those cases when a burst was fired, the first AP was analyzed. **(D)** Distribution of the different event types of SWR-coupled intracellular events for each recorded neuron (control n = 24 cells, post-SE n = 12 cells, same data set as in Fig. 3), depicted as in Fig. 5D. The suprathreshold events, i.e., APs and bursts were pooled. **(E)** Distribution of the % of neurons that can generate APs/bursts, EPSPs or IPSPs during SWRs. **(F)** Quantification of suprathreshold events during SWRs as APs or bursts. *** p < 0.001.

**Figure 7.**
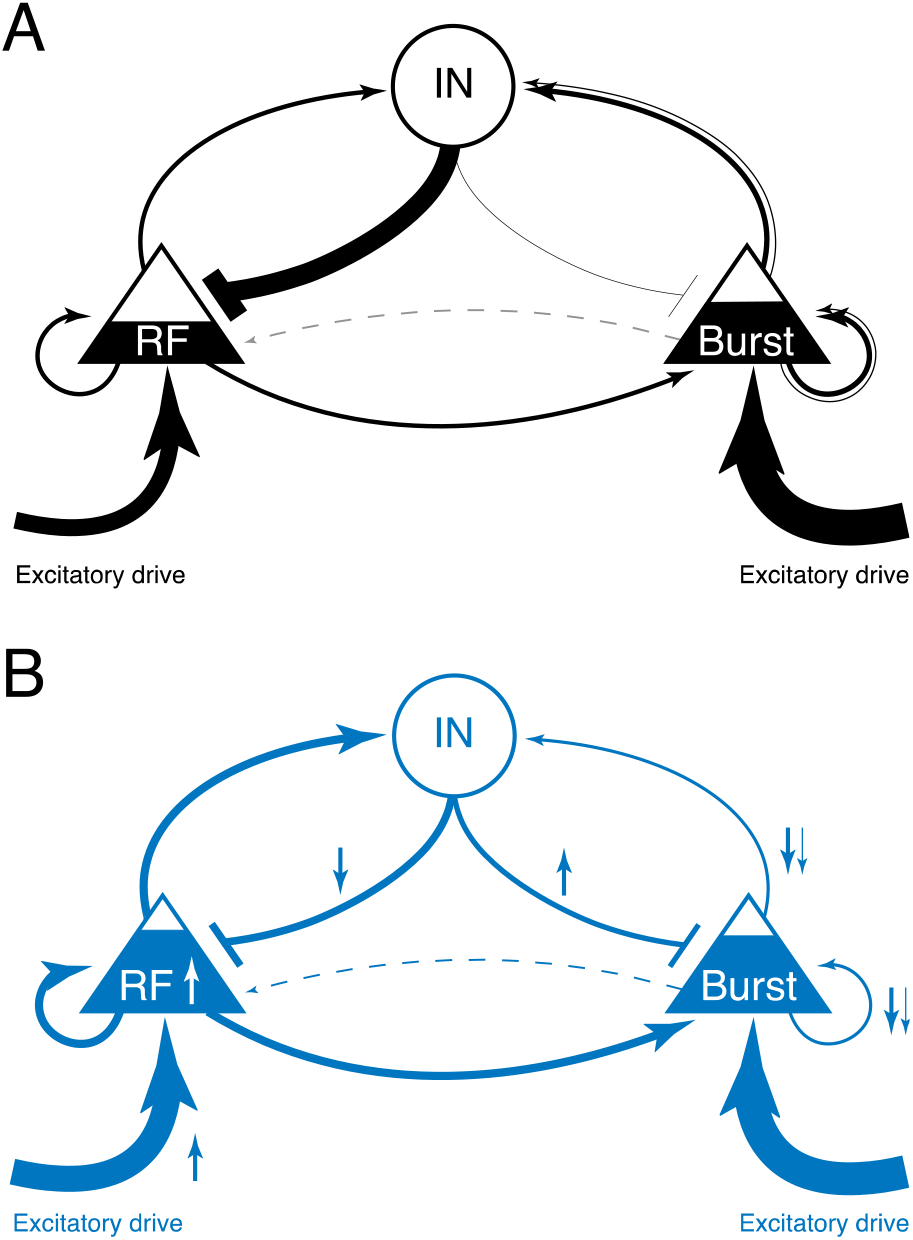
Altered functional connectivity in the subicular microcircuit during SWRs. **(A)** Scheme of a simple microcircuit containing RFs, burster PCs (Burst) and interneurons (IN) in the control condition (modified from Böhm et al., 2015). Thickness of arrow lines between cell types indicate the proportion of inputs (IPSPs, EPSPs) as well as outputs (APs) corresponding to the data of Fig. 5D, 6D and 6F. The second arrow outgoing from the burster PC displays the proportion of generated burst events during SWRs. Filling level (related to the vertical axis only) of RF and burster cell triangles indicate the percentage of neurons firing APs during SWRs. **(B)** Same microcircuit scheme for post-SE neurons. Small arrows aside represent the statistically significant changes compared to control.

**Table 2.**
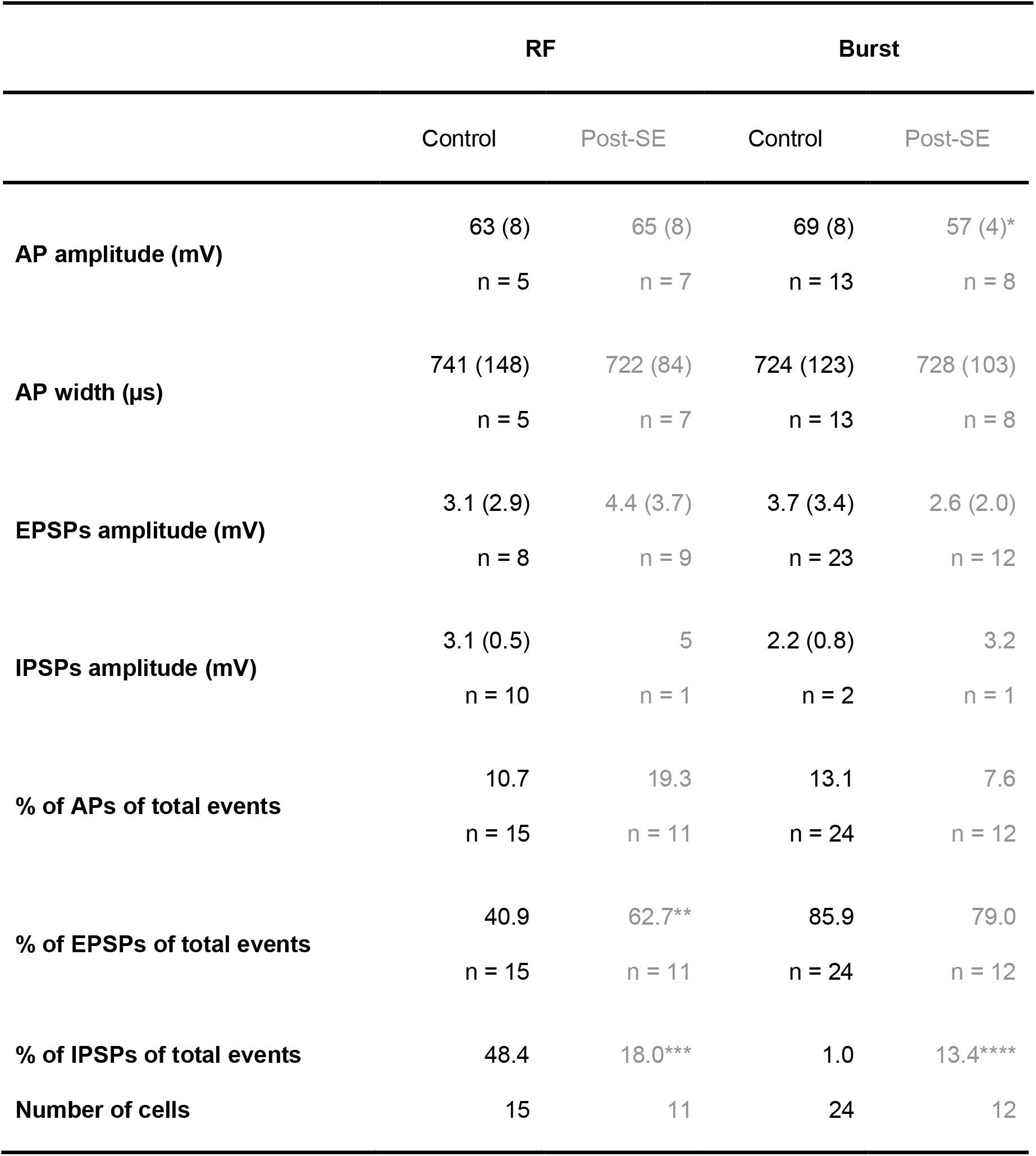
Properties and proportions of subthreshold intracellular postsynaptic potentials as well as APs during subicular SWRs. All data is presented as median (IQR) with n being the number of neurons per group. Significances are tested between control and post-SE groups in RF and burster neurons, respectively. MW, * p = 0.007, Chi-square, ** p = 0.002, *** p < 0.001, **** p = 0.002. EPSPs = excitatory postsynaptic potentials, IPSPs = inhibitory postsynaptic potentials.

|  | RF |  | Burst |  |
| --- | --- | --- | --- | --- |
|  | Control | Post-SE | Control | Post-SE |
| **AP amplitude (mV)** | 63 (8)<br>n = 5 | 65 (8)<br>n = 7 | 69 (8)<br>n = 13 | 57 (4)*<br>n = 8 |
| **AP width (μs)** | 741 (148)<br>n = 5 | 722 (84)<br>n = 7 | 724 (123)<br>n = 13 | 728 (103)<br>n = 8 |
| **EPSPs amplitude (mV)** | 3.1 (2.9)<br>n = 8 | 4.4 (3.7)<br>n = 9 | 3.7 (3.4)<br>n = 23 | 2.6 (2.0)<br>n = 12 |
| **IPSPs amplitude (mV)** | 3.1 (0.5)<br>n = 10 | 5<br>n = 1 | 2.2 (0.8)<br>n = 2 | 3.2<br>n = 1 |
| **% of APs of total events** | 10.7<br>n = 15 | 19.3<br>n = 11 | 13.1<br>n = 24 | 7.6<br>n = 12 |
| **% of EPSPs of total events** | 40.9<br>n = 15 | 62.7**<br>n = 11 | 85.9<br>n = 24 | 79.0<br>n = 12 |
| **% of IPSPs of total events** | 48.4 | 18.0*** | 1.0 | 13.4**** |
| **Number of cells** | 15 | 11 | 24 | 12 |

### Burster neurons fire preferentially single APs rather than bursts during SWRs in post-SE mice

In view of our observation of an increased number of ripples per SPW post-SE, we were particularly interested in the behavior of burster neurons during spontaneously occurring SWRs (Fig. 6A-B). Burster neurons presented besides EPSPs, IPSPs, and single APs also bursts of APs in a fraction of SWRs (Fig. 6B). As with RF neurons, the median AP widths as well as EPSPs and IPSPs amplitudes were similar for the control and post-SE groups in burster neurons (Table 2). However, like the depolarizing step-evoked APs (Fig. 4G), the amplitudes of SWR-associated APs were significantly smaller in post-SE mice (57 (5) mV) compared to controls (69 (8) mV, Mann-Whitney-U test (MW) p = 0.005, Fig. 6C), suggesting that changes in intrinsic excitability of burster neurons are also relevant during network activity without artificial stimulation. Regarding amount of inputs to control burster cells, we observed that only 1 % of all events (in all burster cells) were inhibitory (Fig. 6D) and that 13% of burster neurons received inhibitory inputs during SWRs (Fig. 6E, 7A). While the inhibitory input was weaker than in RF neurons, the excitatory drive was twice as frequent: EPSPs constituted 86% of all events (Fig. 6D) and all burster neurons generated EPSPs during SWRs (Fig. 6E). Burster neurons produced a similar amount of suprathreshold events (13%) compared to RF neurons (11%), which include single APs as well as bursts of APs. However, similar to previously reported data, we observed that the proportion of burster neurons generating APs during SWRs was higher (54%, Fig. 6E) compared to RF neurons (33%, Fig. 5E, Böhm et al., 2015; Eller et al., 2015). While in burster neurons from post-SE mice the proportions of EPSPs and APs did not change, inhibitory events increased from 1 to 13% compared to controls (Chi-square p = 0.002, Fig. 6D, 7B). Moreover, the proportions of in- and outputs did not change in post-SE animals: 17% of neurons displayed IPSPs, 92% EPSPs and 67% fired one AP or a burst during SWRs. Importantly, however, bursts of APs occurred during SWRs in 33% of control neurons while this number decreased to 17% in post-SE (segment ‘AP / Burst’ in Fig. 6E). With regards to the overall event distribution, in control mice 26% of all suprathreshold events during SWRs were bursts of APs, whereas single APs comprised the other 74% (Fig. 6F, double line in 7A). By contrast, the ratio of bursts to single APs changed significantly in post-SE mice, with only 2% being bursts and 98% single APs (Fig. 6F, 7B, Chi-square p < 0.001), potentially due to the increased inhibitory and reduced excitatory drive. To summarize, we observed in controls that burster neurons received predominantly excitatory drive and, during SWRs, half of the burster neurons fired APs, including bursts in a quarter of the cases. In contrast, post-SE burster neurons received more inhibitory drive and fired almost only single APs rather than bursts during subicular SWRs.

## Discussion

In this study, we revealed that an altered cellular contribution may underlie hyperexcitable features in subicular sharp wave-ripples after status epilepticus. We find that after SE 1) SPWs decrease in rate but show a larger amplitude, 2) ripple and unit number is increased with a reduced proportion of memory-formation-associated long-duration ripples, 3) ripples and units build up earlier potentially due to a shifted timing of postsynaptic and action potentials in pyramidal cells, 4) RF neurons receive less inhibitory and more excitatory inputs and consequently fire more often APs during SWRs, while 5) burster neurons receive more inhibition and are more likely to generate only single APs than bursts during SWRs. These modified cellular contributions and hyperexcitable SWRs in the subiculum are likely to impact memory consolidation as well as retrieval and may contribute to epilepsy-associated cognitive dysfunctions due to the strategic important role of the subiculum as a gate to the information storing neocortex.

### Altered SWRs in epilepsy models

Epilepsy develops after status epilepticus in ~20-40% of patients (Sculier et al., 2018) and is frequently associated with cognitive comorbidities like learning and memory dysfunction (Helmstaedter and Elger, 2009; Henin et al., 2021; Hermann et al., 2008). To study the impact of SE-induced epilepsy on memory-associated SWRs, we used the intrahippocampal kainate model (Lévesque and Avoli, 2013). Our *in vivo* recordings confirmed the induction of SE in all animals and chronic seizures in 67% (6 of 9) of animals. Even though we cannot rule out the possibility that the remaining three animals did not develop epilepsy, it is very likely that we missed those seizures in our relatively sparse recordings (1-2 hrs/1-3 weeks) since seizures occur only twice a day (Cavalheiro et al., 1982) and previous studies with more frequent recordings have confirmed electrophysiologically and histologically the development of chronic epilepsy in all observed animals (Bouilleret et al., 1999; Dugladze et al., 2007).

While high-frequency oscillations and epileptiform discharges have been previously characterized in the subiculum of epilepsy patients (Alvarado-Rojas et al., 2015; Cohen et al., 2002), here for the firsttime we demonstrate changes in subicular SWRs from rodents after SE. We describe underlying cellular mechanisms regarding the altered contribution of distinct pyramidal neuronal subtypes to SWRs in kainate-treated animals. Besides, several studies have reported altered SWR rates and ripple frequencies in the epileptic hippocampal CA1 region. Generally, in CA1 the SWR rate decreased in those models (Cheah et al., 2021, 2019; Gelinas et al., 2016; Marchionni et al., 2019), which is in accordance with our observations from the subiculum. However, increases in SWR rates both in hippocampal and extrahippocampal areas in animals developing chronic epilepsy upon intrahippocampal kainate injections have also been reported (Li et al., 2018). Similarly, both increases (Gelinas et al., 2016; Marchionni et al., 2019) and decreases (Cheah et al., 2019; Gelinas et al., 2016) in the ripple frequencies have been shown, while primarily an increased and less-organized CA1 pyramidal cell firing was observed in the intra-amygdalar and intraperitoneal kainate-induced temporal lobe epilepsy models (Foffani et al., 2007; Marchionni et al., 2019; Valero et al., 2017). Altogether, the alterations in SWRs during epileptogenesis are complex and might reflect the various pathologic alterations of the SWR-generating regions, such as CA3 and CA1, compensatory homeostatic changes as well as distinct frequencies of recurrent seizures and subsequent neuronal impairments. Yet, a common characteristic is the increase and desynchronization of neuronal firing after epileptogenesis, strongly implying that SWR-mediated memory engrams are distorted by additional, less precisely timed neuronal activity. Our study focuses for the first time on the combined cellular and network changes of SWRs in the hippocampal gatekeeper region subiculum in an epilepsy model and shows hyperexcitable SWRs with an underlying differential and desynchronized cellular contribution.

### SWRs in the subiculum show features of hyperexcitability and memory impairment

*In vitro* recordings in hippocampal slices from post-SE mice revealed intact propagation of SWRs from CA3 via CA1 to the subiculum. However, the rate of subicular SPWs was reduced, indicating either a diminished generation in CA3, potentially due to a CA3 pyramidal cell loss in the kainate model (Lévesque and Avoli, 2013), or a reduced generation of SWRs in the subiculum itself, which acts as a secondary generator of SWRs (Imbrosci et al., 2021; Norimoto et al., 2013). Reduced SWR rates have been previously reported in context of interictal epileptiform discharges (IEDs) in rodent CA1 (Gelinas et al., 2016) and human hippocampus (Henin et al., 2021) and were associated with poorer performance in a memory task. Indeed, we observed a reduction of a potential marker for memory performance, i.e., the long-ripple duration (Fernández-Ruiz et al., 2019), in post-SE mice, thus supporting the hypothesis that altered SWRs in chronic epilepsy may contribute to poor memory performance.

By contrast, features associated with hyperexcitability and reduced GABAergic inhibition in the neural network (Buzsáki, 2015), such as the amplitude of the subicular SPWs, were increased, whereas the duration of the sharp wave, representing the length of neuronal replay events (Davidson et al., 2009), remained unchanged. This points towards a stronger contribution of local excitatory neurons, most probably RF neurons, due to a hyperexcitable or disinhibited network. However, the amplitude increase was subtle, far below the magnitude of interictal events (Karlócai et al., 2014). The observed hyperexcitable SWRs are therefore referring to actual SWRs as opposed to a replacement of SWRs by IED previously reported for CA1 (Karlócai et al., 2014). Other than previous reports in CA3 and CA1, we failed to observe increased ripple frequencies, within the scope of our ripple bandpass filter (120-300 Hz, Bragin et al., 1999a; Foffani et al., 2007; Ibarz et al., 2010; Jefferys et al., 2012; Jiruska et al., 2017) in post-SE mice. It could be that faster ripples are for example restricted to presubicular hippocampal areas or are absent in our post-SE epilepsy model potentially due to restriction of the ictogenic focus to the dorsal hippocampus (Bragin et al., 1999; Dugladze et al., 2007). Yet, we did see an increased number of ripples per SPW that may either support the hypothesis of a network hyperexcitability or result from an intensified subicular generation of ripples. Furthermore, ripple and unit amplitudes were larger in SWRs from epileptic mice indicating an altered timing such as a more selective synchrony of neuronal firing or an altered contribution of neuronal subtypes, respectively.

### Regular firing neurons contribute to “hyperexcitable SWRs”

Our observation of a higher number of units per SPW is in line with increased activity of RF neurons during SWRs. The increased activity of RF neurons was on one hand associated with a delayed and decreased GABAergic drive, possibly reflecting loss of inhibitory input (Knopp et al., 2008). On the other hand, excitatory drive built up earlier and was significantly increased in frequency. Both seemingly antagonistic phenomena were potentially recruiting an increased proportion of RF neurons to actively fire APs during SWRs in post-SE mice. Increased ripple number and amplitude may thus result from an augmented contribution of disinhibited RF neurons in ripple generation. Besides, we observed an early buildup of ripple and unit activity relative to the SPW trough, which temporally matched most notably the AP firing of RF neurons in post-SE mice. Such early buildup is typical for fast ripples and interictal discharges observed in a previous study in CA1 (Bragin et al., 1999). This early activity buildup might lead by means of recurrent activation of subicular interneurons to an increased inhibitory tone onto burster neurons (Böhm et al., 2015), which is in line with their reduced bursting and potentially the overall delay of ripples relative to the SPW trough post-SE. Because RF neurons project mostly to brain areas involved in retrieval of non-spatial memory, such as context and object-related memory (Böhm et al., 2018), their contribution to information encoding and transfer during SWRs is smaller (Böhm et al., 2018, 2015; Eller et al., 2015; Maslarova et al., 2015). Therefore, their activation seems to be an epiphenomenon of pathologic wiring in the hyperexcitable subicular network (Knopp et al., 2008, 2005) and may actually lead to missense SWRs with incorrect content. Whereas this hypothesis needs to be confirmed in behavioral experiments, some analogy can be seen to a mouse model of schizophrenia (calcineurin KO, characterized by increased synaptic strength), where SWRs recorded from the hippocampus *in vivo* were similarly characterized by an increased number of ripples and units, but the firing sequences of place cells during SWRs no longer corresponded to those during exploratory behavior (Suh et al., 2013).

### Failed Information transfer by inhibited burster neurons

Another major pathologic aspect of subicular post-SE SWRs was the altered contribution of burster neurons. Usually, burster neurons participate actively in SWRs during propagation of hippocampal spatial information to output structures involved in spatial processing (Böhm et al., 2018; Nitzan et al., 2020; Simonnet and Brecht, 2019). This feature is emphasized by the fact that a burst of APs was shown to cause a modification in synaptic strength and to better encode spatial information than a single AP (Lisman, 1997; Simonnet and Brecht, 2019). Yet, in our recordings subicular burster neurons received more inhibitory inputs and the ratio of bursts versus single APs generated during SWRs was dramatically reduced. These observations were also reflected in the intrinsic properties of post-SE burster neurons, since their firing frequency and AP amplitude declined, indicating either an increased inhibitory drive (de la Prida, 2003), increased potassium and/or decreased sodium conductances post-SE. Downregulated excitability of burster neurons might also explain why we did not observe an increased ripple frequency (within the frequency range of physiological ripples) in our recordings as a sign of hyperexcitability, as faster ripples can reflect in-phase synchronous bursting (>300 Hz) of individual burster cells or alternating out-of-phase firing of more than one cell at lower frequencies (100-200 Hz, Ibarz et al., 2010). Indeed, burster neurons in the CA1 from chronically epileptic rodents developed hyperexcitable features due to altered expression of sodium and calcium channels (Becker et al., 2008; Chen et al., 2011), consistent with the generation of fast ripples and their major role in the propagation of epileptic discharges (Hao et al., 2021). However, the present study in the subiculum did not reveal such hyperexcitability in burster neurons. Moreover, a pilocarpine model of epilepsy demonstrated that the number of burster neurons was reduced and that, similar to our observations, the inhibition onto burster neurons was increased (Knopp et al., 2008, 2005). Since the subiculum is the main output gateway of the hippocampus, stronger inhibitory control of subicular burster neurons may prevent seizure generalization (de la Prida, 2003) to downstream areas in temporal lobe epilepsy. Yet, this happens at the cost of impaired information encoding and transfer.

## Conclusion

Here, we show that several SWR features previously linked to memory performance are disrupted in subicular SWRs of mice that underwent kainate-induced status epilepticus. These changes are associated with hyperexcitable attributes and excitation/inhibition imbalances, consequently leading to dominant regular firing as well as silent burster neurons. Furthermore, the time shift of units and ripples relative to the SPW trough implies that the information content carried by SWRs might be altered with consequences for cognitive processes such as memory consolidation. Thus, we suggest that the information content is altered in two different ways at the cellular level, which might be the object of further studies: hyperexcitable features are gained due to hyperactivation of RF neurons, whereas missing features are gone due to loss of function of burster neurons. Direct targeting of these pathological SWR features by, e.g., optogenetically closed-loop silencing of RF or stimulation of burster neurons (Buzsáki et al., 2015; Paz et al., 2013) may in the future present an opportunity to recover memory performance disturbed by epilepsy.

## Materials and Methods

### Experimental license

Animal experiments were performed in accordance with the guidelines of the European Communities Council and approved by the regional authority (LaGeSO Berlin: G0355/10 and T0212/08).

### Stereotactic implantation

Young adult C57BL/6 mice (~ 10 weeks) were deeply anesthetized using isoflurane (1-3% in 100% O2). Mice were then head-fixed in a stereotactic frame and the scalp was exposed. After drilling four holes at the designated locations (Fig S1A), mice were stereotactically implanted with a custom-made assembly made of an injection-cannula (26 gauge, P1 Technologies, Roanoke, VA, USA) and a Teflon-coated platinum-iridium wire (⍉ = 33 μm, Science Products, Hofheim, Germany) into the right dorsal hippocampal area (rHippo) CA1 [coordinates, distance from bregma: anterior - posterior (AnPo) = −2 mm; medial - lateral (ML) = −1.8 mm; dorsal - ventral (DV) = −1.4 mm for cannula and −1.5 mm for wire; Paxinos and Franklin, 2004]. LFP recordings in CA1 were performed therewith in the proximity of the injection site. For ECoG recordings miniature stainless-steel screws were implanted above the right frontal lobe (rFront: AnPo = 1.5 mm, ML = −1.5 mm), the left parietal lobe (lPariet: AnPo = −2 mm, ML = 1.8 mm) and for reference above the right cerebellum. Implanted electrodes were secured on the skull with dental acrylic resin (UNIFAST Trad, GC America Inc., Alsip, IL, USA). Screws were soldered to insulated silver wires (AG-10T, Science Products, Hofheim, Germany), which together with the platinum electrode were separately soldered to an electrode interface board (EIB-8, Neuralynx, Bozeman, MT, USA). After surgery, mice were housed individually and left to recover for at least 2 days.

### Induction of status epilepticus

For induction of SE, 50 nl of 20 mM kainic acid (‘kainate’, Sigma-Aldrich, Taufkirchen Germany, solved in 0.9% NaCl) were injected over 10 min into right dorsal hippocampal area CA1. This was performed in freely moving animals by using a 0.5 μl microsyringe (7000 Series, Hamilton Company, Reno, NV, USA) that was connected via a connector-assembly (C313C/SPC, P1 Technologies, Roanoke, VA, USA) to the implanted cannula. After infusion the syringe was left in place for an additional 2 min to limit reflux along the cannula track.

### In vivo data acquisition and analysis

Recordings were collected for 1 - 2 hours in freely moving mice before injection, right after injection during the ongoing SE and on average four times at 1 - 9 weeks after SE. SE was determined by *in vivo* recordings and visually detected tonic-clonic activity. During recordings, electrophysiological signals were pre-amplified by a tethered headstage (HS-8-CNR-MDR50; Neuralynx) that was connected to the EIB-8 interface board. Signals were amplified, band-pass filtered (1 Hz - 10 kHz) and continuously acquired at 30 kHz (Digital Lynx, Neuralynx). For seizure detection data were stored in the smr file format (Spike2, Cambridge Electronic Design Limited, Milton, UK) and inspected manually for epileptic activity.

### Slice preparation and solutions

Implanted mice (6 - 9 weeks post induction of SE) and age-matched controls were decapitated under isoflurane anesthesia (1-3% isoflurane, in 70% N_2_O, 30% O_2_) and the rapidly removed brain was transferred into ice-cold carbogenated (95% O_2_, 5% CO_2_) artificial cerebrospinal fluid (aCSF) containing in mM: NaCl 129, NaHCOs 21, KCl 3, CaCl_2_ 1.6, MgSO_4_ 1.8, NaH_2_PO_4_ 1.25, and glucose 10 with pH adjusted to 7.4 and osmolarity to 300 ± 3 mOsmo/l. Horizontal hippocampal-entorhinal cortex slices, 400 μm thick, were prepared on a vibratome (LEICA VT1200 S, Leica Biosystems, Wetzlar, Germany) as described previously (Behrens et al., 2005) and immediately transferred to an interface chamber, perfused with carbogenated aCSF at 36 ± 0.5°C (flow rate: ~1.8 ml/min). Slices were left to recover for at least 2 hours before recording.

### In vitro recordings and data acquisition

Extracellular field potentials were recorded in an interface chamber using glass microelectrodes filled with 154 mM NaCl (5 - 10 MΩ). Electrodes were placed in the pyramidal layer of the subiculum. In some recordings additional electrodes were placed in the pyramidal layers of area CA3 and CA1 to detect the propagation pattern of SWRs. LFPs were amplified 200 times with a custom-built amplifier and low-pass filtered at 3 kHz. To correlate intracellular signals and extracellular SWRs, intracellular recordings were performed in parallel using sharp microelectrodes (70–90 MΩ), pulled from borosilicate glass (o.d. 1.2 mm) and filled with 1.8 M potassium acetate. Intracellular signals were amplified with a SEC 05L amplifier (NPI Instruments, Tamm, Germany). All signals were digitized at 5-10 kHz (1401 interface, CED, Cambridge, UK) and recorded with Spike2 software.

### SWR detection and analysis

Data from LFP recordings were analyzed offline and underwent first a DC removal (time constant 0.5 s) in Spike2. SWR data were then analyzed using custom-written scripts in Matlab (The MathWorks Inc., Natick, MA, USA). For sharp wave (SPW) detection data were lowpass filtered (cut frequency 80 Hz, 8th order, symmetrical Butterworth filter) and the offset, i.e., the median of the entire lowpass signal, was subtracted from the lowpass data. SPWs were detected, when a threshold (multiple (0.9-3, mean: control: 1.83, post-SE 1.98, Mann-Whitney-U test p = 0.45) of twice the standard deviation (SD) of the lowpass data mean, i.e., the cross level) crossed the lowpass signal. Subsets of data were manually reviewed and thresholds for automated detection were adjusted if necessary. Adjacent events (inter-event interval < 15 ms) were merged and events with durations below 5 ms or above 200 ms were discarded. For each data segment below the cross level, i.e., the detected SPW, the local minimum, i.e., the SPW trough, was identified for preparing the calculation of the SPW amplitude and counting the number of SPWs per minute. For determining correct SPW durations and amplitudes the threshold was refined for each SPW event by adding 15 ms before (start) and after (end) the cross levels. Out of these two 15 ms segments the median was calculated. The actual beginning of a SPW was redefined when this median was crossed between start and three datapoints after the cross level. The end of a SPW was redefined, when the median crossed a second time in the range of 3 datapoints before the second cross level and the end of the SPW. This was set as the beginning and end of the SPW event, respectively. The distance between the beginning and end point was determined as the SPW duration. The SPW amplitude was calculated by subtracting the individual median of the 15 ms segments from the minimum of the SPW event. For ripple and unit detection data were bandpass filtered (120-300 and 600-4000 Hz, respectively) by an 8th order symmetrical Butterworth filter (He et al., 2010). Ripples and units were detected in a range of 100 ms before and after the SPW through. Thresholds for detecting ripples and units were 5 and 6 times the SD of the bandpass filtered data, respectively. Averaged frequencies for each event were calculated between consecutive minima. Amplitudes were determined by averaging the amplitudes from the preceding and succeeding maxima for each minimum. The durations between the first and last ripple or unit through, respectively, per SPW were calculated. All SPWs, ripples and units, respectively, were grouped (Fernández-Ruiz et al., 2019; Imbrosci et al., 2021) according to their treatment (control vs. post-SE).

### Electrophysiological characterization

The intrinsic properties of neurons were characterized in intracellular recordings as described previously (Maslarova et al., 2015). Briefly, we used depolarizing and hyperpolarizing steps of increasing intensity (from ±70 pA to ± 560 pA in 70 pA intervals, 300 ms duration). Only cells with a membrane potential more negative than −50 mV and input resistance (defined as the voltage-current ratio at the second half of the hyperpolarizing step, after the initial sag had subsided) higher than 20 MΩ were accepted for analysis. APs were detected with a threshold of −30 mV. The first inter-spike interval for each current step was inverted for estimation of the instantaneous firing rate as a function of the current. Neurons were categorized as bursters if they fired a burst of APs upon depolarization with an instantaneous frequency of at least 120 Hz and the AP burst was nested over a depolarizing envelope, followed by an after-hyperpolarization of at least 2 mV. The sag and rebound (reb) ratios were calculated as described previously (de la Prida et al., 2003): Sag ratio = (Vstep-Vsag) / Vstep, and Reb ratio = (Vreb-Vsag) / Vreb. Vstep is the maximum voltage offset caused by the hyperpolarizing step; Vsag is the mean voltage during the second half of the hyperpolarization step, when the depolarization caused by the hyperpolarization-activated current (Ih) is stable; and Vreb is the voltage at maximum rebound.

### Statistical analyses

Data are presented as single data points and the associated box plots presenting the median and the Q25-Q75. In the text, data are stated as median and interquartile range (IQR). Statistical comparison was conducted using the unpaired two-tailed student’s t-test (t-test) for normally distributed numerical data, the Mann-Whitney-U test (MW) for non-parametric numerical data and the Kolmogorov-Smirnov test (KS) for probability distributions (IgorPro, Wavemetrics, Lake Oswego, OR, USA). For comparisons of dependent data, the paired t-test was used (Sigma Plot, Systat Software, Inc., San Jose, CA, USA). Categorical data was compared using the two-tailed Chi-square test (GraphPad Software, San Diego, CA, USA). P-values (p) < 0.05 were considered as statistically significant.

## Acknowledgements

We thank the late Uwe Heinemann for help on the conception of this project and for his mentorship. We thank Alexey Ponomarenko for help on the *in vivo* recordings, Therese Alich for help on the experimental setup and Heinz Beck as well as Istvan Mody for providing experimental resources and comments on an earlier version of this project. We thank Jens Eilers for comments on the manuscript.

## Author Contributions

KL and AM designed the study. KL, ZJK, SS and AM performed the experiments. JOH created the Matlab analysis script. KL and AM analyzed the data and wrote the manuscript.

