## Supplemental Figure 1 for "Status epilepticus induces chronic silencing of burster and dominance of regular firing neurons during sharp wave-ripples in the mouse subiculum"

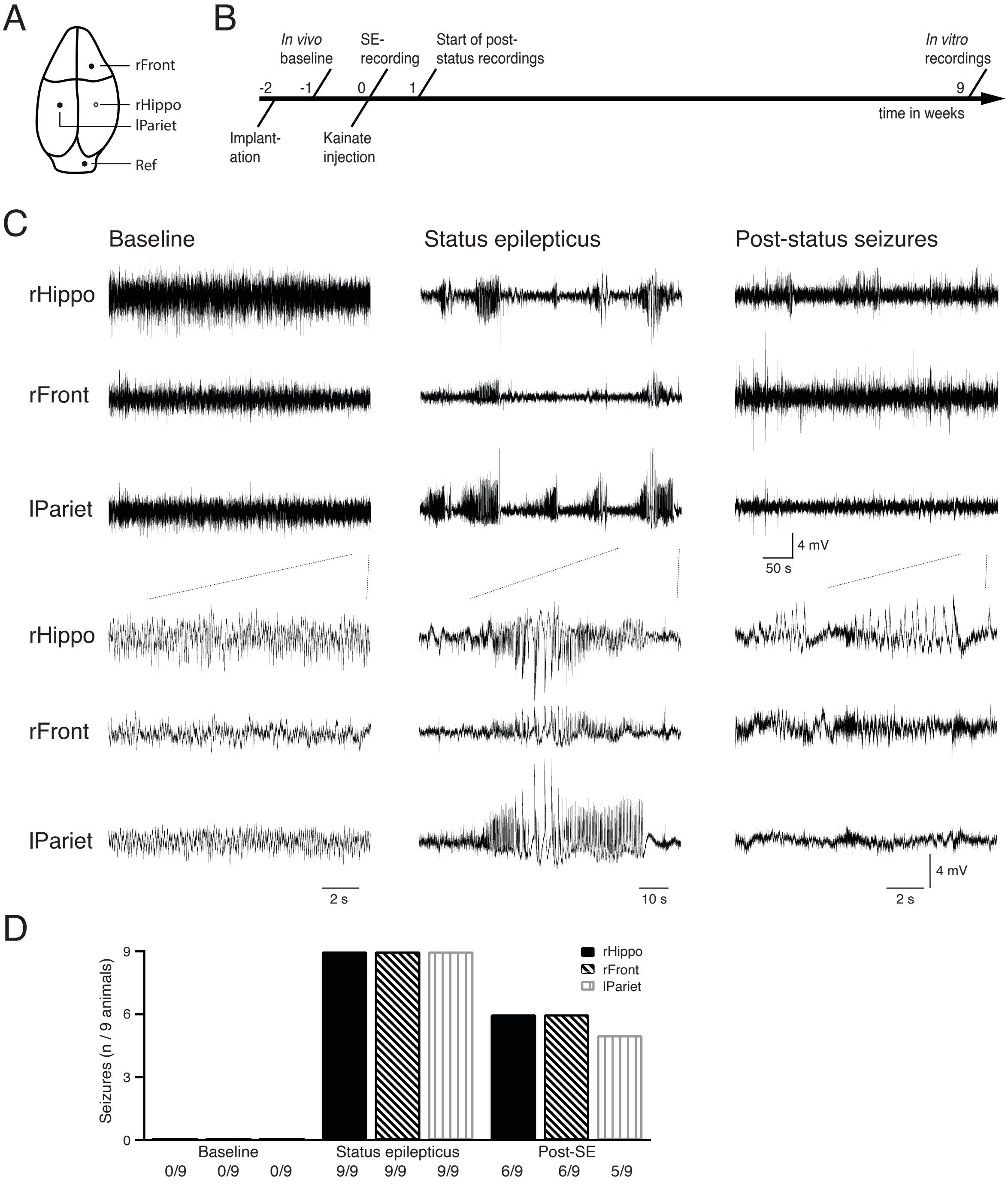


**Supplementary Figure S1** Injection of kainic acid (kainate) in the dorsal hippocampal area CA1 induces a status epilepticus and chronic seizures. **(A)** Schematic mouse skull displaying the recording sides of the LFP recording at the right hippocampal area CA1 (rHippo, open circle), the electrocorticographic (ECoG) recordings (closed circle) from the right frontal lobe (rFront), the left parietal lobe (lPariet) as well as the right cerebellum used as reference (Ref). **(B)** Timeline of experiments showing the electrode implantations two weeks and the baseline recording within one week before the kainate-induced status epilepticus. Several recordings of two hours each were obtained up to nine weeks post-status epilepticus. **(C)** Example LFPs from the three recording sites with zoom-ins (three lower panels) from pre-status baseline (left), status epilepticus (middle) and eight weeks post-status seizures (right) from the same animal. **(D)** Bar graph showing the number of animals (total n = 9) that presented with seizures at pre-status baseline, during SE and post-status seizures according to the recording side.
